# Reliable identification of protein-protein interactions by crosslinking mass spectrometry

**DOI:** 10.1101/2020.05.25.114256

**Authors:** Swantje Lenz, Ludwig R. Sinn, Francis J. O’Reilly, Lutz Fischer, Fritz Wegner, Juri Rappsilber

## Abstract

Crosslinking mass spectrometry is widening its scope from structural analyzes of purified multi-protein complexes towards systems-wide analyzes of protein-protein interactions. Assessing the error in these large datasets is currently a challenge. Using a controlled large-scale analysis of *Escherichia coli* cell lysate, we demonstrate a reliable false-discovery rate estimation procedure for protein-protein interactions identified by crosslinking mass spectrometry.

---

Crosslinking mass spectrometry has become a key technology for structural biology by providing distance restraints on purified multi-protein complexes^1^. Proteins can also be crosslinked in complex mixtures. Charting protein-protein interactions (PPIs) in cell lysates, organelles or even whole cells has, therefore, become the next frontier for this technique^2–13^. To avoid reporting large numbers of spurious PPIs, it is important that the false discovery rate (FDR) of the reported interactions is reliably estimated. Doing this correctly is a challenge for the field^12–15^ and to date, a consensus has not yet emerged (**Table S1**).

The target-decoy search strategy is a reliable approach for error estimation in proteomics that has been adapted for crosslinking^16–19^. It assumes that the rate of hits to the ‘decoy’ database is an estimator of false positives (or type II error rate). Here, with experimental validation, we show that two additional considerations need to be addressed for it to correctly estimate errors in crosslinking-based PPI screens. Firstly, crosslinks involving two different proteins (heteromeric crosslinks) must be considered separately during FDR estimation from those involving the same protein (self-links, including homomeric crosslinks) due to their inherently different search spaces (**Fig. 1a, Fig. S1**). Since most possible crosslinks in the database are heteromeric, most false positives in the total (self and heteromeric) set of crosslinks will be heteromeric. Controlling FDR in the total set, and then selecting only heteromeric matches, thus enriches for false positives (**Fig. S1**). Secondly, crosslink-spectra matches (CSMs) must be merged into PPIs for reliable FDR estimation of PPIs (**Fig. 1b**).

**Figure 1:**
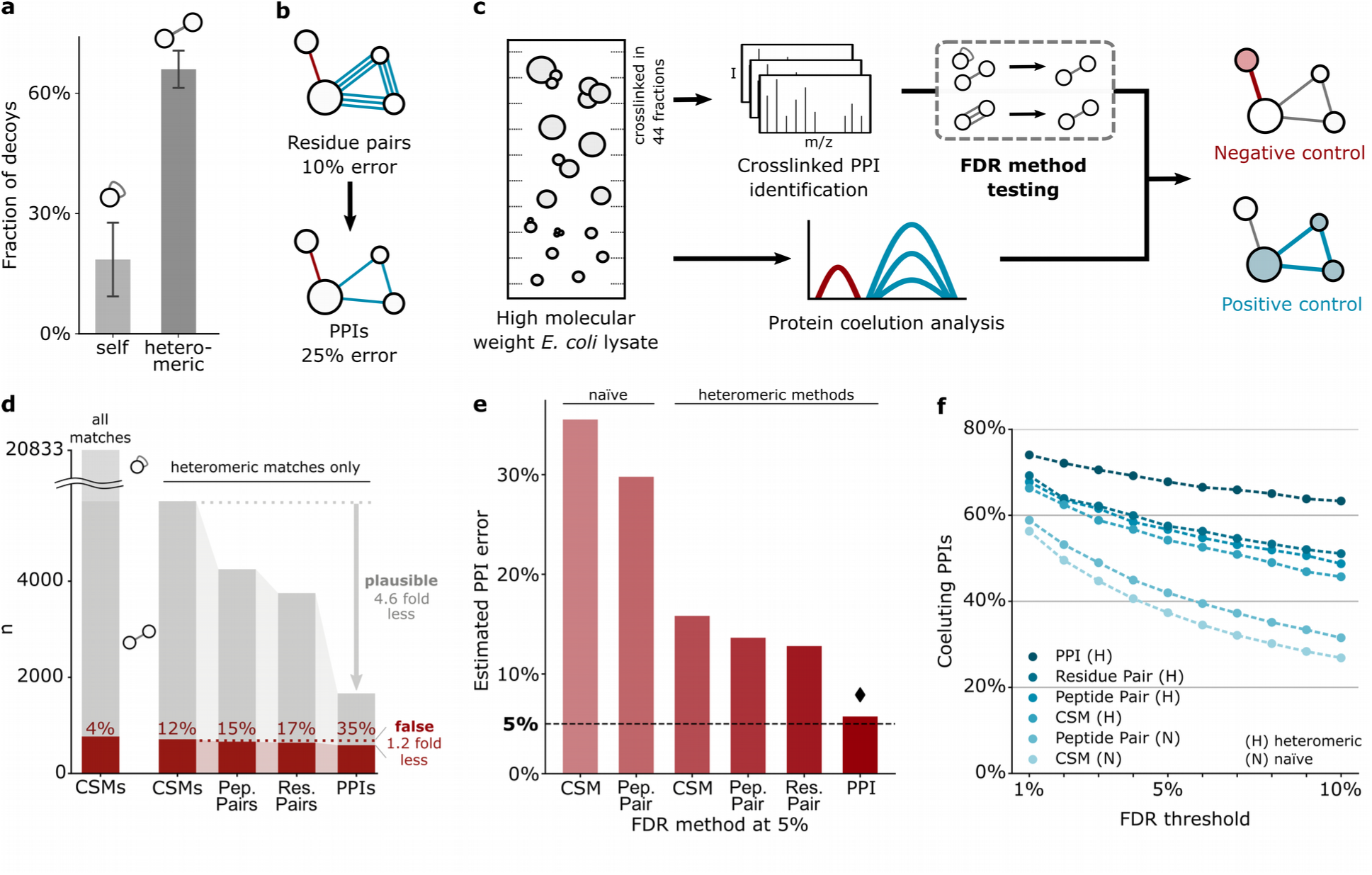
Heteromeric PPI-FDR method for reliable identification of crosslinked protein-protein interactions. a) Fraction of decoys in 10 random picks of 100 self and 100 heteromeric PPIs from the raw search output. Error bars show standard deviation from the mean. b) Schematic showing error increase when merging crosslinked residue pairs to PPIs. c) Experimental workflow. *E. coli* lysate was separated and crosslinked in individual high molecular weight fractions, pooled again to simulate a complex mixture, and analyzed by mass spectrometry. Quantitative proteomics of uncrosslinked fractions provided protein coelution data. d) False identifications as a function of merging heteromeric CSMs passing a ‘naïve’ CSM-FDR of 5% (BS3 data, for DSSO data see **Fig. S4a**). When merging crosslink data from CSMs to PPIs, the number of identifications decreases and the fraction of false identifications increases. e) PPI error resulting from a 5% FDR threshold of published FDR approaches (**Table S1**). Diamond denotes the method leading to the PPI error closest to 5%. Mean of BS3 and DSSO data is shown (separated in **Fig. S4b**). f) Fraction of protein pairs with similar elution profiles (correlation coefficient > 0.5) among the PPIs passing a given FDR threshold, applying different published FDR approaches (**Table S1**). Average of BS3 and DSSO data (separated in **Fig. S4c**).

To test different approaches for PPI-FDR estimation, we designed a controlled large-scale crosslinking study of lysate from the model organism *E. coli* (**Fig. 1c**). Lysate was fractionated by size exclusion chromatography and the proteins in each of 44 high molecular weight fractions were analyzed by quantitative proteomics to generate protein elution profiles. In parallel, aliquots from each fraction were crosslinked with bis-sulfosuccinimidyl suberate (BS3) or the MS-cleavable disuccinimidyl sulfoxide (DSSO), respectively. These fractions were pooled for each crosslinker to simulate the complexity of a bacterial cell. Proteins that were not found to be eluting together in the same fraction could not have been crosslinked, so we used these ‘non-crosslinkable’ protein pairs as an independent validation of the estimated FDR (**Fig. S2**). Following protein digestion, the crosslinked peptides were fractionated extensively (2 × 90 fractions, 32.5 days of mass-spectrometric acquisition) to generate a substantial dataset.

We first searched against a database comprising all *E. coli* proteins, including those not detected in our sample. We identified 20,833 (5655 heteromeric) unique CSMs for BS3 and 22,296 (6923 heteromeric) unique CSMs for DSSO at a ‘naïve’ 5% CSM-level FDR (not distinguishing self and heteromeric links). We chose 5% to have sufficient false identifications for precise FDR estimation at all levels. Our control revealed that 4% of these CSMs reported false PPIs, i.e. interactions between proteins either found in different fractions or not identified in our sample by proteomics (**Fig. 1d, Fig. S3**). Merging CSMs into heteromeric PPIs (as in **Fig. 1b**) revealed that a ‘naïve’ 5% CSM-FDR led to 36% of PPIs being false (35% for BS3, **Fig. 1d**; 36% for DSSO, **Fig. S3**). In an alternative control that doubled the database by adding human protein sequences (entrapment database), the PPI error with a calculated ‘naïve’ 5% CSM-FDR reached 47% (**Fig. S3**). Of note, 87% of false PPIs involved proteins that lacked self-links (100% in the entrapment control) (**Fig. S4**). These proteins in PPIs lacking self-links had a lower median abundance than all proteins in the sample suggesting that they are random matches (**Fig. S4**). In contrast, proteins with self-links had significantly higher abundance.

The inflated error of the ‘naïve’ CSM-FDR was in part caused by not assessing heteromeric matches separately from self-matches. Such separation decreased false PPIs substantially (35% to 16% and 36% to 15%, for BS3 and DSSO, respectively). However, the error remained three times higher than the targeted 5%, which linked to error propagation. CSMs rarely corroborated each other in false PPIs while plausible PPIs were supported by multiple CSMs (average 1.2 versus 4.6 CSMs) (**Fig. 1b,d**). This was most pronounced when merging unique residue pairs into PPIs. Error control at lower information levels therefore leads to large proportions of reported PPIs being false (**Fig. 1e, Fig. S5**). In contrast, controlling the error at PPI level by merging CSMs for each PPI prior to FDR estimation gave more reliable results: 5.7% false PPIs when applying 5% PPI-FDR (**Fig. 1e**). This also applies to other FDR thresholds and the entrapment control which indicated 4% error when applying 5% PPI-FDR (**Fig. S6**). As a positive control, we evaluated the proportion of PPIs that were supported by correlation of protein coelution profiles (**Fig. 1f, Fig. S7**) or interaction evidence from the STRING database (**Fig. S8**). The fraction of ‘known’ PPIs was highest when using our heteromeric PPI-FDR and the proportion decreased when raising the FDR threshold, as expected (**Fig. 1f, Fig. S8**). These concepts were implemented in our open source FDR estimation software tool, xiFDR v2.0, which is search software independent.

To focus on a high-quality subset of PPIs in the *E. coli* lysate, we applied a 1% heteromeric PPI-FDR cutoff, yielding 590 PPIs involving 308 proteins (**Fig. 2a, Table S2**), connected with a total of 2539 residues pairs. 366 (62%) of these PPIs are connected by more than one residue pair (**Fig. S9**). 11% of the proteins found in PPIs were flagged as potential false matches by having no self-links. Their abundance is nevertheless significantly higher than random, and they tend to be small proteins and thus generally difficult to observe (**Fig. S9**). We found 63% (370) in the STRING database (**Fig. 2b**). 98% (576) were found to be eluting in a fraction together and 68% had similar elution profiles (correlation coefficient > 0.5), suggesting that they form stable complexes (**Table S3**). Ribosomal proteins displayed a complex elution pattern, presumably due to the presence of assembly intermediates, although many of the proteins that were found crosslinked to the ribosome are known interactors (26 of 53).

**Figure 2:**
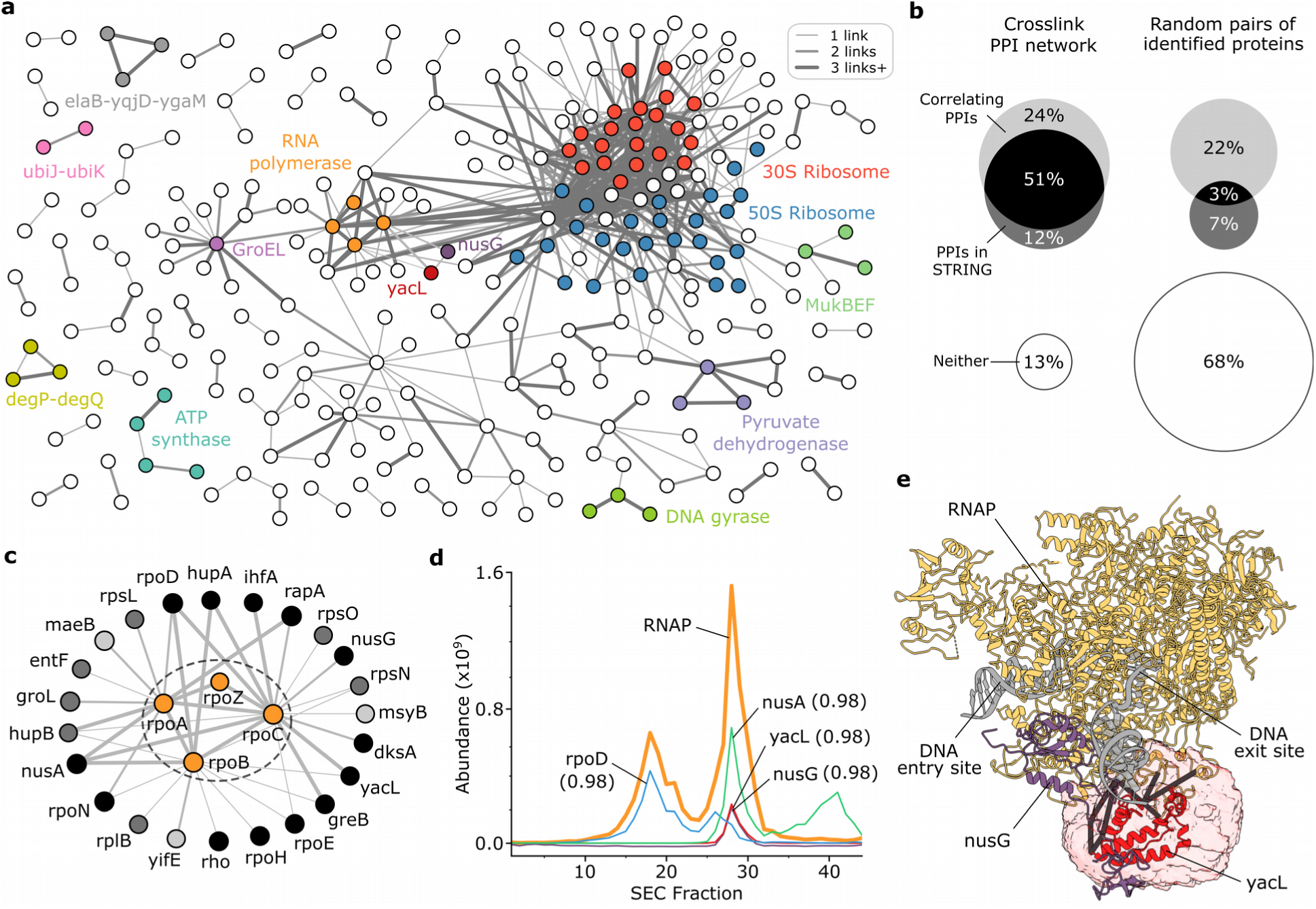
Heteromeric PPI-FDR leads to high fidelity PPI network in *E. coli* lysate. a) CLMS-derived PPI network of soluble high molecular weight *E. coli* proteome. Selected proteins and protein complexes are highlighted. The proteins aceA and tnaA were removed for clarity. b) PPI network overlap with STRING database and coelution data (correlation coefficient > 0.5) compared to random pairs of identified proteins. c) PPI subnetwork of RNAP. Color scheme for RNAP binders according to categories from panel b. d) Elution traces of RNAP (average abundance of its subunits) and selected RNAP binders with their minimal elution correlation coefficient to any RNAP constituent. e) RNAP with bound NusG (PDB 5MS0) and the region of CLMS-defined accessible interaction space for YacL (I-TASSER model) highlighted.

The crosslink-based PPI network (**Fig. 2a**) included 289 protein pairs with highly similar coelution (correlation coefficient > 0.8). The majority of these were known interactions including complexes like ATP synthase, pyruvate dehydrogenase, MukBEF or DNA gyrase (**Fig. S10 - S13**). The data confirmed binding of acyl carrier protein to MukBEF, and of YacG, to DNA gyrase (**Fig. S12, S13**). In addition, 130 PPIs with highly similar coelution were not yet experimentally confirmed for *E. coli* K12, though 55 of these had a STRING entry based on other evidence. Novel interactions included those between the small ribosomal regulators ElaB, YgaM and YqjD, the periplasmic endoproteases DegP and DegQ, the ubiquinone biosynthesis accessory factors UbiK and UbiJ (**Fig. S14**), as well as GroEL and potential substrates (**Fig. S15**).

RNA polymerase (RNAP) linked to 23 proteins (**Fig. 2c**). Previous interaction evidence was available for 20 of these, including the transcription factors RpoD and GreB, and the transcriptional regulators NusG, NusA and RapA; all crosslinks are in agreement with previously suggested binding sites (**Fig. S16**). YacL, a protein of unknown function that was found to be associated with RNAP in pull-down experiments^20^, coeluted with RNAP (correlation of 0.988) with an abundance comparable to NusG (**Fig. 2d**). Crosslinks localized YacL on RNAP next to NusG at the DNA exit site (**Fig. 2e**).

Correctly controlled error is an important element of any discovery-based technology. Here, we demonstrated a reliable procedure to estimate the FDR of protein-protein interactions in crosslinking mass spectrometry. We showed that separating self and heteromeric crosslinks and merging CSMs into PPIs prior to FDR estimation are required to report PPIs with reliable confidence. Note that when aiming at distance restraints for modelling of protein complexes, residue pair FDR should be considered. In any case the FDR threshold can be chosen to meet the stringency required by the study (**Fig. S17**). Crosslinking mass spectrometry can now bridge the gap between structural studies and systems biology by reliably revealing topologies of PPIs in their native environments.

## Acknowledgements

We would like to thank Richard Scheltema, Alexander Leitner, Andrea Sinz, Michael Hoopmann, Marc Wilkins, Fan Liu, Henning Urlaub, and Dermot Harnett for comments on the manuscript. The work was funded by the Deutsche Forschungsgemeinschaft (DFG, German Research Foundation) under Germany’s Excellence Strategy – EXC 2008 – 390540038 – UniSysCat and grant no. 426290502 and by the Wellcome Trust through a Senior Research Fellowship to JR (103139). The Wellcome Centre for Cell Biology is supported by core funding from the Wellcome Trust (203149).

## Author Contributions

F.O., L.S., S.L. and J.R. designed the experiments; L.S., F.W. and F.O. prepared the samples. L.S., S.L. and L.F. collected and processed CLMS data; L.F. designed and implemented xiFDR software; S.L., L.S., F.O., and J.R. prepared figures and wrote the manuscript with input from all authors.

