## Supplemental Tables, Figures & Methods for "Reliable identification of protein-protein interactions by crosslinking mass spectrometry"

**This PDF file includes:**

Materials and Methods

Table S1

Figures S1-S17

**Other Supplementary Materials for this manuscript include the following:**

Table S2: Detected heteromeric PPIs at 1% FDR

Table S3: Protein quantitation from SEC fractionation

Table S4: All plausible PPIs based on SEC fractionation

Table S5: Self and heteromeric unique residue pairs of detected PPIs

### Materials & Methods

#### Materials

Unless otherwise stated, reagents were purchased in the highest quality available from Sigma (now Merck, Darmstadt, Germany). Empore 3M C-18-Material for LC-MS sample cleanup was from Sigma (St. Louis, MO, USA), glycerol from Carl Roth (Karlsruhe, Germany). The BS3 (bis(sulfosuccinimidyl) suberate) crosslinker was supplied by Pierce Biotechnology (Thermo Fisher Scientific, Waltham, MA, USA) and the DSSO (disuccinimidyl sulfoxide) crosslinker from Cayman Chemical (Ann Arbor, MI, USA).

#### Biomass production

A single clone of *Escherichia coli* K12 strain (BW25113 purchased from DSMZ, Germany; https://www.dsmz.de/) grown on Agar plates was selected for inoculation of lysogeny broth (LB)-media. A preculture aliquot was used to start fermentation in a Biostat A plus bioreactor (Sartorius, Göttingen, Germany) in LB medium with 0.5% (w/v) glucose and at 37°C. The pH and dissolved oxygen were monitored and adjusted by the addition of sodium hydroxide/phosphoric acid or stir speed control, respectively. Overall growth was monitored by optical density measurements at 600 nm. When the culture reached an optical density of 10, the fermentation was stopped and the culture rapidly cooled in stirred ice water followed by harvesting the biomass by centrifugation at 5000 g, 4°C for 15 min. Cell pellets were stored at -80 °C after washing with PBS and snap-freezing in liquid nitrogen.

**Cell lysis and high-molecular-weight proteome fractionation by size exclusion chromatography (SEC)**

Cell pellets were suspended at 0.2 g wet-mass per ml in ice-cold lysis buffer (50 mM Hepes pH 7.2 at RT, 50 mM KCl, 10 mM NaCl, 1.5 mM $MgCl_2$, 5% (v/v) glycerol, 1 mM dithiothreitol (DTT), spatula tip of chicken egg white lysozyme (Sigma, St. Louis, MO, USA)). Cells were lysed by sonication on ice. Prior to sonication cOmplete EDTA-free protease-inhibitor cocktail (Roche, Basel, Switzerland) was added according to the manufacturer's instructions. After sonication, 125 units of Benzonase (Merck, Darmstadt, Germany) were added. Subsequently, the lysate was cleared of cellular debris by centrifugation for 15 min at 4°C and 15.000 g. DTT was added again to 2 mM. This cleared lysate was subjected to ultracentrifugation using a 70 Ti fixed-angle rotor for 1 h at 106.000 g and 4°C. Then, the supernatant was concentrated using ultrafiltration with Amicon spin filters (15 kDa molecular weight cut-off) to reach a total protein concentration of 10 mg/ml, as judged by microBCA assay (Thermo Fisher Scientific, Waltham, MA, USA). Aggregates were removed by centrifugation for 5 min at 16,900 g and 4°C. 1 mg of soluble high molecular weight proteome was loaded onto a BioSep SEC-S4000 column (600 × 7.8 mm, pore size 500 Å, particle size 5 µm, Phenomenex, CA, USA) and fractionated at 200 µl/min flow rate and 4°C while collecting fractions of 200 µl over the separation range from ~3 MDa to 150 kDa (as judged by Gel filtration calibration kit (HMW), GE Healthcare).

**Sample preparation for LC-MS protein identification with non-crosslinked SEC fraction aliquots**

For protein identification, aliquots (40 µl) of each fraction were precipitated by adding four volumes of cold acetone followed by an incubation at -20°C overnight. Pellets were collected by centrifugation and supernatants discarded. Protein pellets were air-dried and subsequently solubilized using 6 M Urea, 2 M Thiourea, 100 mM ABC (ammonium bicarbonate). Derivatization was accomplished by incubating for 30 min at RT with 10 mM DTT followed by 20 mM IAA (iodoacetamide) for 30 min in the dark at RT, respectively. Proteases were added to the samples: LysC (1:100 (m/m)) for 4.5 h at 37°C, followed by diluting 1:5 with 100 mM ABC and continued with with trypsin (1:25 (m/m)) at 37°C for 16 h. The reactions were stopped by adding TFA (trifluoroacetic acid) to a pH of 2-3. Subsequently, sample clean-up following the Stage-tip protocol was performed and samples were stored at -20°C until LC-MS acquisition.

**LC-MS protein identification with non-crosslinked SEC fraction aliquots**

Protein elution profiles via LC-MS were acquired using a Q Exactive HF mass spectrometer (Thermo Fisher Scientific, Bremen, Germany) coupled to an Ultimate 3000 RSLC nano system (Dionex, Thermo Fisher Scientific, Sunnyvale, USA). 0.1% (v/v) formic acid and 80% (v/v) acetonitrile, 0.1% (v/v) formic acid served as mobile phases A and B, respectively. Samples were loaded in 1.6% acetonitrile, 0.1% formic acid on a Easy-Spray column (C18, 50 cm, 75 µm ID, 2 µm particle size, 100 Å pore size) operated at 45 °C and running with 300 nl/min flow. Peptides were eluted with the following gradient: 2 to 6% buffer B in 1 min, 6 to 10% B in 2 min, 10 to 30%B in 37 min, 30 to 35% in 5 min followed by 35 to 45%B in 2 min. Then, the column was set to washing conditions within 1.5 min to 90% buffer B and flushed for another 5 min. For the mass spectrometer the following settings were used: for MS1 scans resolution was 120,000, AGC (automatic gain control) target $3*10^6$, maximum injection time 50 ms, scan range from 350 to 1600 m/z. The ten most abundant precursor ions with z = 2-6, passing the peptide match filter (“preferred”) were selected for HCD (higher-energy collisional dissociation) fragmentation employing stepped normalized collision energies (29 +/-2). The quadrupole isolation window was set to 1.6 m/z. Minimum AGC target was $2.5*10^4$, maximum injection time was 80 ms. Fragment ion scans were recorded with a resolution of 15,000, ACG target set to $1*10^5$, scanning with a fixed first mass of 100 m/z. Dynamic exclusion was enabled for 30 s after a single count and included isotopes. Each LC-MS acquisition took 75 minutes.

**Quantitative proteomics database search**

Raw data from bottoms-up proteomics experiments were processed using MaxQuant[21] version 1.6.0.16 operated under default settings (fully tryptic digestion with two missed cleavages maximum; up to five variable modifications (oxidised methionine and acetylated protein N-termini, MS1 match tolerance 20 ppm (first search) / 4.5 ppm (main search), MS/MS match tolerance 20 ppm) per peptide; carbamidomethylation of cysteine set as fixed modification; 1% PSM and protein group FDR). Each SEC fraction was treated as an individual experiment. Quantitation by iBAQ[22] requiring a minimum of two peptides (unique+razor) and matching between runs were enabled. The database used was the Uniprot curated reference proteome UP000000625 with two unreviewed entries removed summing to a total 4350 proteins (downloaded on 04/08/2019).

**Protein crosslinking, digestion and sample cleanup of SEC fractions**

The remaining parts of the SEC fractions (160 µl, see above) were split into two 75 µl aliquots, for the two crosslinking reactions, and adjusted to 97.5 µl with 1x SEC-Running buffer. Crosslinker stock solutions were prepared freshly at 30 mM in water free DMF. Crosslinking of the fractions was initiated by quickly mixing each sample with 2.5 µl crosslinker stock to a final concentration of 0.75 mM crosslinker. The crosslinking reaction was incubated for 2 h on ice before quenching with ABC at 50 mM and further incubation for 30 min on ice. The crosslinked samples were acetone-precipitated at -20°C overnight (see above). Protein was solubilized in 6 M Urea, 2 M Thiourea, 100 mM ABC. For sample reduction and alkylation, 10 mM DTT for 30 min at RT and 20 mM IAA for 30 min at RT, dark were employed. For sample proteolysis, LysC was added at 1:100 (m/m) ratio and incubated for 4 h at 37°C. Upon 1:5 dilution with 100 mM ABC, Trypsin was added to the sample (1:25 (m/m)) and digestion continued for 16 h at 37°C followed by stopping via addition of TFA to a pH of 2-3. The digests were desalted using SPE cartridges following the manufacturer's instruction and eluates dried, aliquoted and stored at -20°C until further use.

**Multidimensional offline fractionation of crosslinked peptide samples**

All crosslinked peptide pools were fractionated using an Äkta pure system (GE Healthcare, Chicago, IL, USA) employing a PolySulfoethyl A SCX column (100 x 2.1 mm, 300 Å, 3 µm) equipped with a guard column of identical stationary phase (10 x 2.0 mm) (PolyLC, Columbia, MD, USA) running at 0.2 ml/min for the first separation dimension. Here, mobile phase A consisted of 10 mM $KH_2PO_4$ pH 3.0, 30% ACN while mobile phase B contained 1 M KCl in addition. The system was kept at 21°C throughout the fractionation. Dried digestion aliquots of 400 ug peptides were dissolved in mobile phase A. Upon injection, peptides were eluted isocratically for 2 min followed by an exponential gradient up to 700 mM KCl with following steps: 12 min to 12.7%, followed by 1-min steps to 14.5, 16.3, 18.8, 23.0, 30.0, 40.0, 70.0% B. Fractions of 200 µl size were collected over the elution range. The same fractions from five replica SCX runs were pooled for desalting using Stage-tips. Dried Stage-tip eluates of each individual SCX fraction were then subjected to the second dimension offline-fractionation by hSAX chromatography. Here, a Dionex IonPac AS-24 hSAX column (250 x 2.0 mm) with a AG-24 guard column (Thermo Fisher Scientific, Dreieich, Germany) were used on Äkta pure system (see above). Mobile phase A consisted of 20 mM Tris*HCl pH 8.0 with mobile phase B containing 1 M NaCl in addition. The system was kept at 15°C for these experiments. Samples were eluted from Stage-tips, dried and resuspended in mobile phase A. Again, peptides were loaded under isocratic conditions for 3 min, and then eluted by an exponential gradient with the following steps 1.8, 3.5, 5.3, 7.1, 9.1, 11.2, 13.5, 16.3, 19.7, 24.1, 30.2, 38.8, 51.5, 70.6, 100% B lasting for one minute each. Fractions of 150 µl size were collected throughout the elution phase. Adjacent fractions were pooled to give ten pools in total (fractions 3-6 / 7-14 / 15-17 / 18-19 / 20-21 / 22-23 / 24-25 / 26-27 / 29-29 / 30-35), that were desalted using Stage-tips.

**LC-MS for crosslink identification**

LC-MS/MS analysis of crosslinked peptides was performed using a Q Exactive HF mass spectrometer (Thermo Fisher Scientific, Bremen, Germany) coupled to an Ultimate 3000 RSLC nano system (Dionex, Thermo Fisher Scientific, Sunnyvale, USA). Mobile phases A and B consisted of 0.1% (v/v) formic acid and 80% (v/v) acetonitrile, 0.1% (v/v) formic acid, respectively. Samples were loaded in 1.6% acetonitrile, 0.1% formic acid on a Easy-Spray column (C18, 50 cm, 75 µm ID, 2 µm particle size, 100 Å pore size) running at 300 nl/min flow and kept at 45 °C. Analytes were eluted with the following gradient: 2 to 7.5% buffer B in 5 min, followed by a linear 80-min gradient of 7.5 to 42.5% and an increase to 50% B over 2.5 min. Then, the column was set to washing conditions within 2.5 min to 95% buffer B and flushed for another 5 min.

The mass spectrometric settings for MS1 scans used were: resolution set to 120,000, AGC of $3*10^6$, maximum injection time of 50 ms, scanning from 400-1450 m/z in profile mode. The ten most intense precursor ions that passed the peptide match filter ("preferred") and with z = 3-6 were isolated using a 1.4 m/z window and fragmented by HCD using in-house optimized stepped normalized collision energies (BS3: 30 +/- 6; DSSO: 24 +/- 6). Fragment ion scans were acquired at a resolution of 60,000, ACG of $5*10^4$, maximum injection time of 120 ms scanning from 200-2000 m/z, underfill ratio set to 1%. Dynamic exclusion was enabled for 30 s (including isotopes). In-source-CID was enabled at 15 eV to minimize gas-phase associated peptides. Each LC-MS run took 120 minutes.

#### Crosslink database search

Raw data from mass spectrometry was processed using msConvert (version 3.0.11729)[23] including denoising (top 20 peaks in 100 m/z bins) and conversion to mgf-file format. Precursor and fragment masses were recalibrated to account for mass shifts during measurement. Obtained peak files were analysed using xiSEARCH 1.6.746[5] with the following settings: MS1/MS2 error tolerances 3 and 5 ppm, allowing up to 2 missing isotope peaks (Lenz et al., 2018), tryptic digestion specificity with up to 2 missed cleavages, carbamidomethylation on cysteine as fixed and oxidation on methionine as variable modification, losses: $–CH_3SOH$ / $–H_2O$ / $–NH_3$, cross-linker BS3 (138.06807 Da linkage mass) or DSSO (158.0037648 Da linkage mass) with variable cross-linker modifications on linear peptides ("BS3-NH2" 155.09463 Da, "BS3-OH" 156.07864 Da, "DSSO-NH2" 175.03031 Da, "DSSO-OH" 176.01433 Da). For samples crosslinked with DSSO, additional loss masses were defined accounting for its cleavability ("A" 54.01056 Da, "S" 103.99320 Da, "T" 85.98264). Crosslink sites for both reagents were allowed for side chains of Lys, Tyr, Ser, Thr and the protein N-terminus.

The same database as for the non-crosslinked samples was used. For the entrapment database control the database was extended by the same number (4,353) of human proteins. For each *E. coli* protein, a human protein similar in Lysine and Arginine content, as well as sequence length was sampled. For the final PPI network, the search database was reduced to only proteins identified in our 44 SEC fractions, to reduce noise in the database. Decoys were generated for all searches, including the entrapment database. For this, protein sequences were reversed and for each decoy protein the enzyme specific amino acids were swapped with their preceding amino acid[21].

**FDR calculation**

Results were filtered prior to FDR to crosslinked peptide matches having a minimum of three matched fragments per peptide, a delta score of 15% of the match score and a peptide length of at least six amino acids. Additionally, identifications ambiguously matching to two proteins or more were removed. FDR was calculated based on decoy matches by xiFDR (version 2.0dev) using the formula[18]:

$$FDR = \frac{TD - DD}{TT}$$

Depending on the experiment, FDR was employed on different result levels (CSM, peptide pair, residue pair or protein pair) with defined thresholds. Scores of higher levels were calculated as described by Fischer and Rappsilber[18]:

$$Score_{higher\ level} = \sqrt{\Sigma\ (Score_{lower\ level})^2}$$

FDR was solely calculated based on that score, no further improvement by other information was done at this point. To account for the improvement of the identification by prefiltering on lower levels[18], the same threshold was employed on each of the lower levels. Self- and heteromeric crosslinks were handled together or separately by enabling / disabling the grouping option.

For the final PPI network, BS3 and DSSO PPIs were separately filtered to 1% heteromeric PPI-FDR. As the scores cutoff differed between the two datasets, the score was normalised to range between 0 and 1 for subsequent joining of the two lists. The combined list was then filtered again to 1% FDR.

**Non-crosslinkable control**

Since crosslinking was performed in individual SEC fractions, proteins without sufficient abundance in the respective fraction are unlikely to produce detectable amounts of crosslinked peptides. For the definition of non-crosslinkable proteins, we determined at which protein abundance (iBAQ) heteromeric peptide pairs are observable. For this, 10% heteromeric peptide pair FDR was calculated on DSSO and BS3 data. For each combination of proteins with a detected peptide pair, the maximum iBAQ of the less abundant protein in each SEC fraction was determined. The cutoff was chosen at the 5-percentile (iBAQ of 4.3E6). Protein pairs for which both proteins never reached that iBAQ value in the same fraction were defined as non-crosslinkable. Therefore, this control does define false interactions between two otherwise plausible proteins. This is different from the human entrapment database which defines only false interactions involving proteins absent from the sample. 544,274 (6% of all possible PPIs) PPIs are defined as plausible (see Table S4), while 8,914,801 (94%) PPIs are non-crosslinkable.

Additionally, the difference in false and plausible search space was taken into account for error estimation. While all matches in the false search space are by definition false, also some matches in the plausible search space will be random. Account for these, the Lysine/Arginine content of proteins in the respective groups was used as an estimate for the number of possible peptides and the error calculated as:

$$Error_{PPI,\,non-crosslinkable} = \frac{n_{false}}{n_{plausible} + n_{false}} \cdot \frac{KR_{plausible} + KR_{false}}{KR_{false}}$$

$$Error_{PPI,\,non-crosslinkable} = \frac{n_{false}}{n_{plausible} + n_{false}} \cdot 1.09$$

where $n_{false}$ is the number of PPIs defined as “non-crosslinkable”, $n_{plausible}$ the number of PPIs that are plausible, both after passing the respective FDR calculation. $KR_{plausible}$ and $KR_{false}$ are the sums of Lysines and Arginines in the proteins of the interactions, respectively. The factor 1.09 is database specific.

**Human entrapment database control calculation**

As a second control, the error of matched PPIs was estimated based on known wrong matches of an entrapment database with human proteins (see above). PPIs were defined as false if one or more proteins in the PPI was a human protein. Additionally, the difference in entrapment and possible search space was taken into account, similar to the approach for the non-crosslinkable control:

$$Error_{PPI,\,entrapment} = \frac{n_{entrapment}}{n_{E.\,coli} + n_{entrapment}} \cdot \frac{KR_{E.\,coli} + KR_{entrapment}}{KR_{entrapment}}$$

$$Error_{PPI,\,entrapment} = \frac{n_{entrapment}}{n_{E.\,coli} + n_{entrapment}} \cdot 1.998$$

As expected from doubling the original database size by adding an entrapment database of equal size, the search space normalisation approximates to 2 (1.998).

**Correlation of protein elution profiles**

Proteins were quantified in each SEC fraction as described above. iBAQ values for each protein were normalized by the maximum of the respective protein over the course of fractionation, leading to normalized abundance values between 1 and 0. For each combination of proteins, elution peaks were detected via the scipy python package (1.4.1). In an elution window of 7 or more fractions, the abundances of the proteins were correlated (Pearson). PPIs with elution profiles with a correlation coefficient > 0.5 were counted as having similar elution profiles. Code was written in python 3.7.

**PPI network comparison with STRING database**

For all *E. coli* K12 proteins identified in quantitative proteomics experiments, interaction evidence from the STRING database v10.5[24] was used (scores ranging from 0 to 1000, retrieved from https://string-db.org on 19th December 2018). PPIs were accounted as known if the STRING combined score was equal or higher than 150. PPIs were defined as lacking experimental evidence when the STRING experimental score was lower than 150. Note that STRING defines 150 as the lowest cutoff in favour of an interaction.

**Plotting protein elution profiles**

For the creation of protein elution profiles, fraction-wise iBAQ intensities for each protein from the MaxQuant search were used (see above), hereinafter referred to as abundance. Individual proteins are represented by their gene names while protein complexes, when shown, are labelled with their respective complex name in the figures. Abundance values for protein complexes were averaged for all components as listed in EcoCyc[25], (retrieved from https://ecocyc.org/ on 25th September 2018). Plots were created in python 3.7 with pandas 0.24.2 using the seaborn 0.9.0 package.

**Protein structural models**

Models of protein complexes with mapped residue pairs (see Table S5) were prepared with xiVIEW[26], python 3.7 with pandas 0.24.2 and ChimeraX 0.92[27].

All structural PPI models were downloaded from the protein data bank (https://www.rcsb.org/): ATP synthase (PDB 5t4O[28]), DNA gyrase (PDB 6RKW[29]), GroEL (PDB 4PKO[30]), RapA (PDB

4S20[31]), GreB (PDB 6RIN[32]), NusG (PDB 5MS0[33]), NusA (PDB 6FLQ[34]) and RpoD (PDB 4ZH3[35])

### Model of RNA polymerase with bound YacL

An I-TASSER[36] model for YacL was generated with default settings based on the Uniprot sequence (see above). DisVis[37] ran under default settings, fixed model was PDB 5MS0 (RNAP, DNA and NusG only), scanning model YacL. Residue pairs of YacL to RNAP and NusG were used as restraints with a minimal distance of 2 Å, maximal distance of 15 Å. The density displayed in Figure 2e corresponds to the accessible interaction space with 5 satisfied restraints. The I-TASSER model was placed for visualisation purposes.

### Data availability

Raw data and MaxQuant outputs from quantitative proteomics experiments were deposited with the ProteomeXchange Consortium partner repository jPOSTrepo under the accession JPST000843 / PXD019004[38].

All raw data, peak lists and search result files from crosslinking experiments were deposited with the ProteomeXchange Consortium partner repository jPOSTrepo under the accession JPST000845 / PXD019120[38].

**Supplementary Table 1:** Recent large scale crosslinking mass spectrometry studies and aspects of their FDR method

| Sample | Search / FDR Software | Self- / heteromeric crosslinks separated for FDR | FDR level | threshold | reference |
|---|---|---|---|---|---|
| Murine synaptosomes | XlinkX / PD | no | CSM | 2% | [7] |
| *Saccharomyces cerevisiae* nucleus | XlinkX / PD | no | CSM | 1% | [9] |
| *Saccharomyces cerevisiae* mitochondria | pLink1 | no | CSM | 1% | [10] |
| Several previously published datasets | pLink2 | yes | CSM | 5% | [4] |
| *Drosophila melanogaster* embryo lysate | MeroX | yes | CSM | 1% | [12] |
| Human cells | Comet | no | Peptide pair | 1% | [2] |
| Human cell lysate | MaxLinker | no | Peptide pair | 1% | [13] |
| Human cell lysate | XlinkX / PD | no | Peptide pair | 1% | [3] |
| *Saccharomyces cerevisiae* mitochondria | Kojak | yes | Peptide pair | 2% | [8] |
| Human cell lysate | xiSEARCH / xiFDR | yes | Residue pair | 5% | [5] |
| Human mitochondria | xiSEARCH / xiFDR | yes | Residue pair | 5% | [6] |
| *Mycoplasma pneumoniae* cells | xiSEARCH / xiFDR | yes | Residue pair & PPI | 5% | [11] |

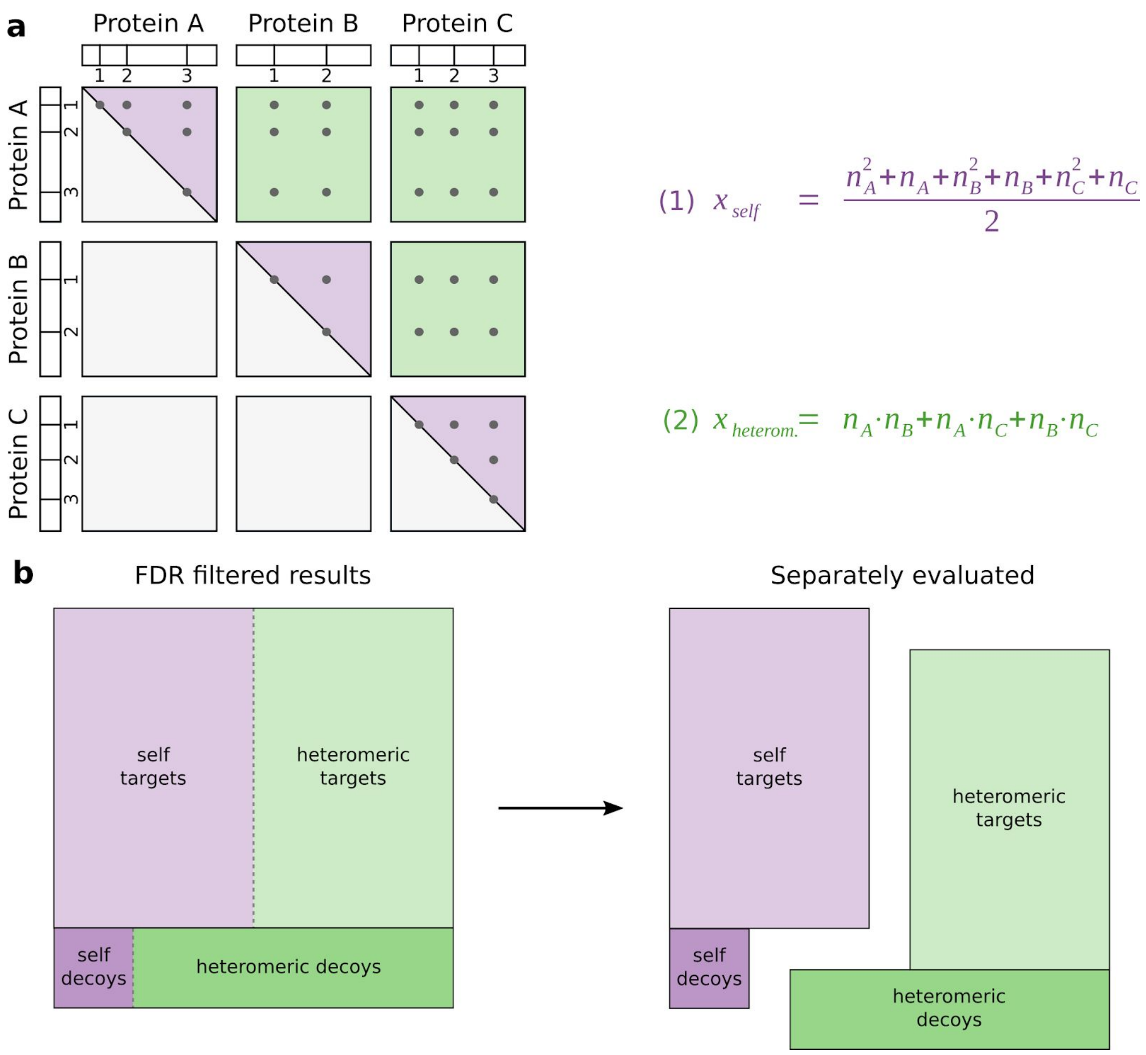


**Supplementary Figure 1: Random spaces for self and heteromeric crosslinks.** Crosslinked peptide spectrum matches consist of two peptides. Due to this, matches can be from the same protein (self) or from two different (heteromeric) proteins. a) For matches within the same protein sequence (with a non-directional crosslinker[19]), a crosslink from A1 to A2 is indistinguishable from A2 to A1. Therefore the number of theoretical possible linksites is defined as (1) (purple triangles). In contrast, heteromeric matches are not symmetrical, and therefore occupy a larger random space which equals to (2) (green squares). b) Illustration of the distribution of decoy matches for self and heteromeric crosslinks. If considered together for FDR calculation, heteromeric decoy matches will be matched more frequently and make up most of the summed decoy matches. If subsequently heteromeric matches are evaluated separately (e.g. for reporting PPIs), their error will be larger than the previously calculated FDR (which would only be correct for the data as a whole).

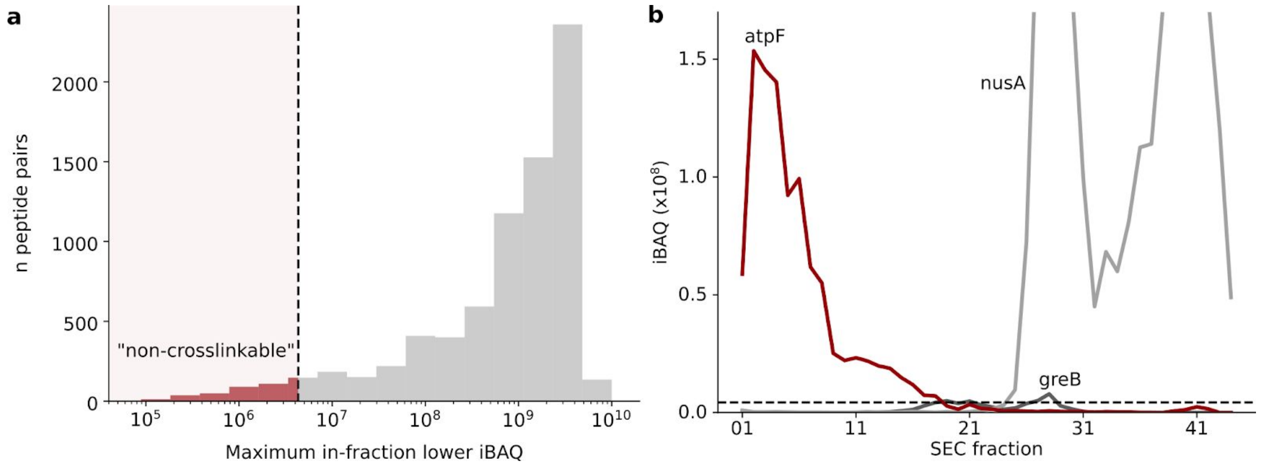


**Supplementary Figure 2: Non-crosslinkable control definition.** a) Distribution of the maximum iBAQ value of the less abundant protein for identified heteromeric peptide pairs. A cutoff for insufficient abundance was chosen at the lower 5-percentile. b) Example of proteins defined either as plausible to crosslink to nusA (greB, dark grey) or “non-crosslinkable” (atpF, red). Although atpF is present in some fractions together with nusA, it is too low abundant in those, therefore the proteins are defined as "non-crosslinkable". In contrast, greB reaches the cutoff in the same fractions with nusA, therefore the two are crosslinkable.

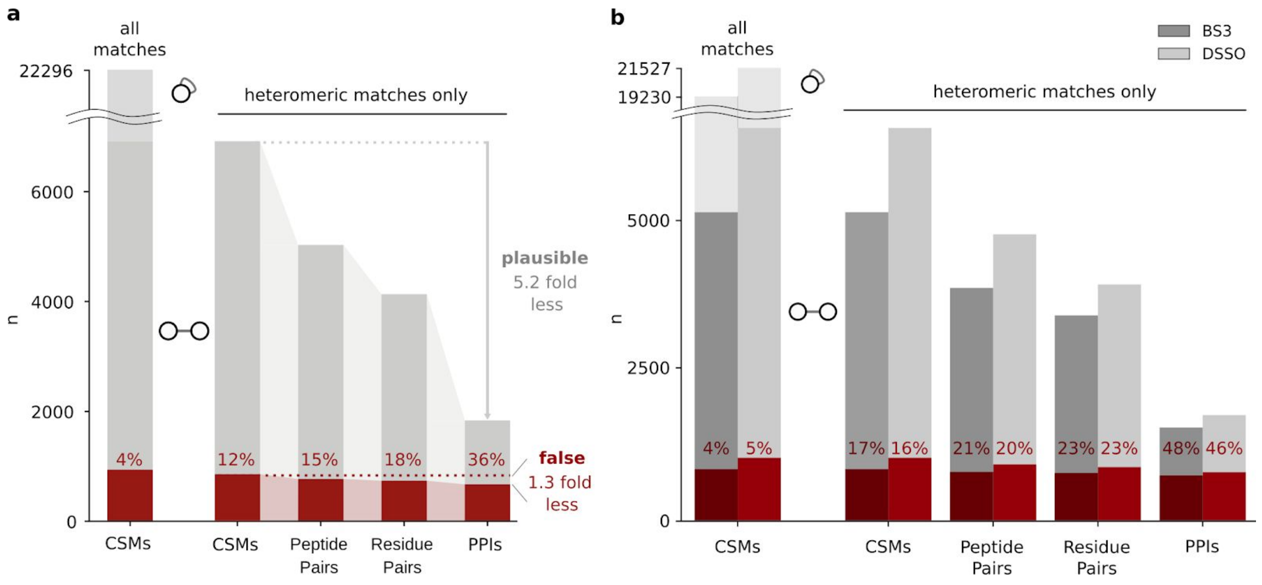


**Supplementary Figure 3: False identifications as a function of merging heteromeric CSMs.** Identifications passing a "naïve" CSM-FDR of 5% for a) DSSO data (corresponding to Fig. 1d) and b) results of the human entrapment database search. The number of false identifications is based on the known false, non-crosslinkable or human, corrected by the respective database sizes (see methods).

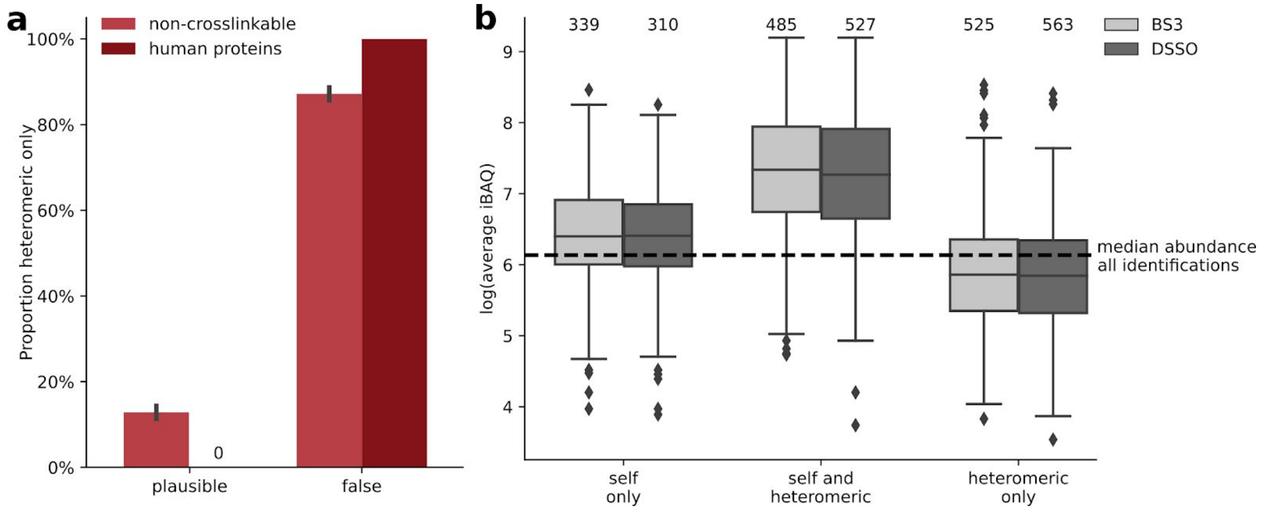


**Supplementary Figure 4: False crosslinks involve proteins lacking self-links.** a) Proportion of PPIs with one or more proteins lacking self-links in the false and plausible PPIs of both negative controls, *E. coli* proteins not found in the same fraction (non-crosslinkable) and human proteins in the entrapment database. b) Proteins found exclusively in heteromeric PPIs had a lower intensity than all identified proteins (median shown by dashed line) and thus a low chance to be detected. In contrast, the median abundance of proteins detected in self-PPIs was 1.9-fold higher than the median of all identified proteins. This increased to 14.8-fold for proteins with both, self-PPIs and heteromeric PPIs (significantly higher abundance than all identified proteins, $p < 0.0001$ using a one-sided Kolmogorov–Smirnov test).

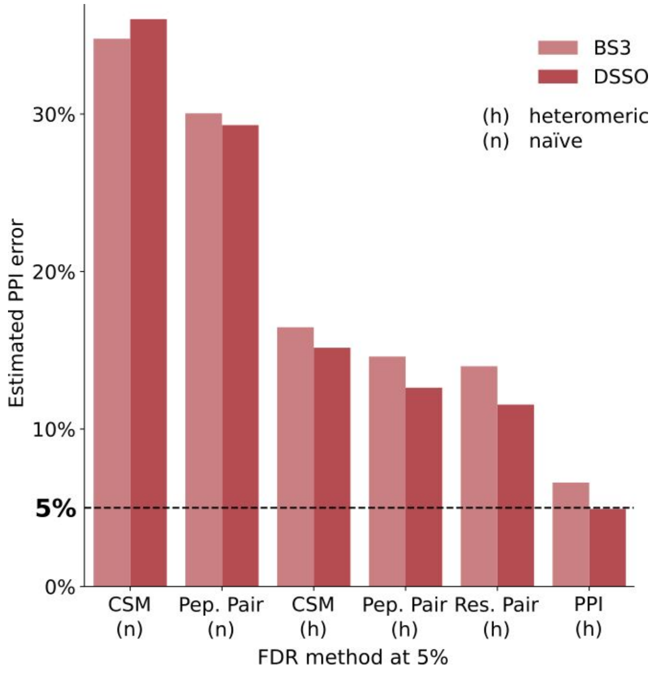


**Supplementary Figure 5: Heteromeric PPI error resulting from a 5% FDR threshold of published FDR approaches (Table S1).**

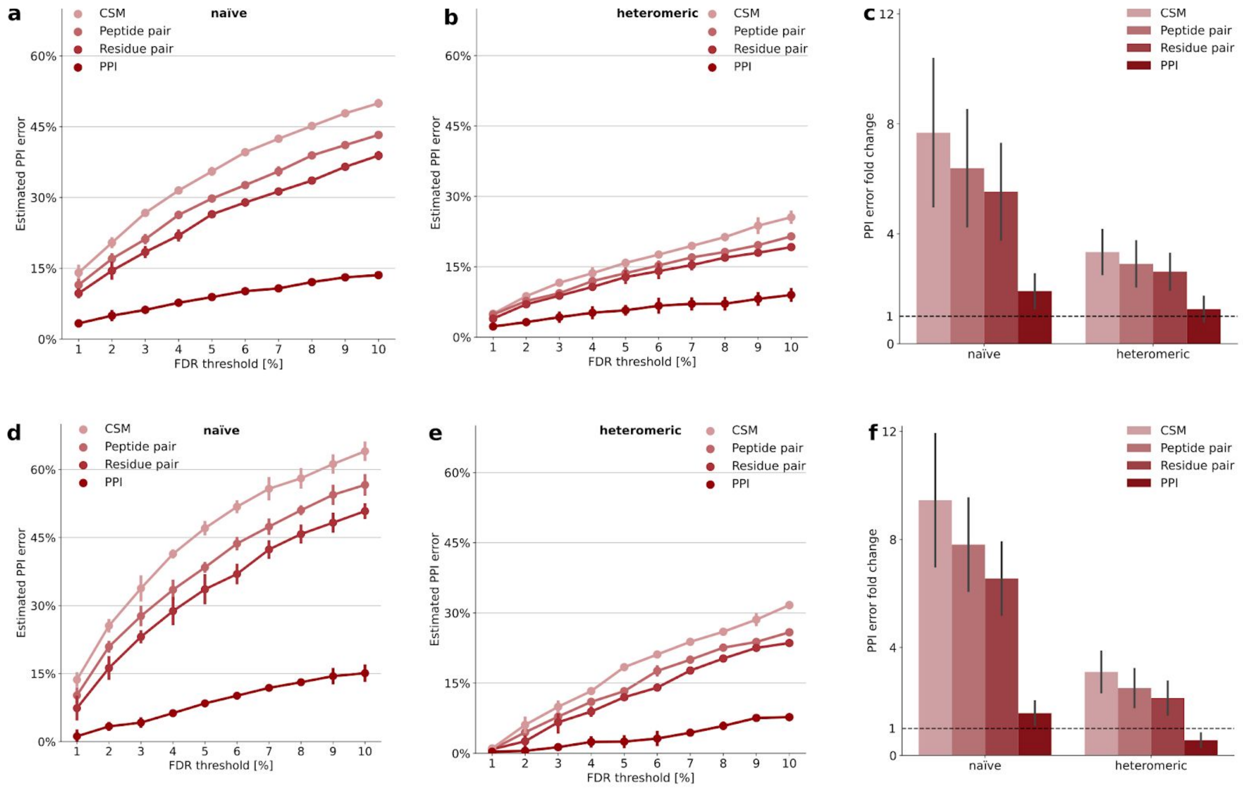


**Supplementary Figure 6: Heteromeric PPI-FDR method applied to non-crosslinkable and entrapment controls, separated by crosslinker.** a-c) Errors of the non-crosslinkable control (see Fig. S2 for definition). Individual errors of different FDR levels for methods calculating FDR on a) all and b) heteromeric crosslinks for the non-crosslinkable control. Error bars represent the standard deviation for BS3 and DSSO datasets. c) Average fold changes (experimental error / desired FDR cutoff) for FDR methods in the non-crosslinkable control. Error bars represent the standard deviation of the individual FDR thresholds for each crosslinker dataset. d, e) Errors of the human entrapment database control. Individual errors of different FDR levels for methods calculating FDR on d) all and e) heteromeric crosslinks for the human control. Error bars represent the standard deviation for BS3 and DSSO datasets. Note that at low FDR thresholds the calculated error is less reliable due to small numbers of false matches. f) Average fold changes (experimental error / desired FDR cutoff) for FDR methods in the human control. Error bars represent the standard deviation of the individual FDR thresholds for each crosslinker dataset.

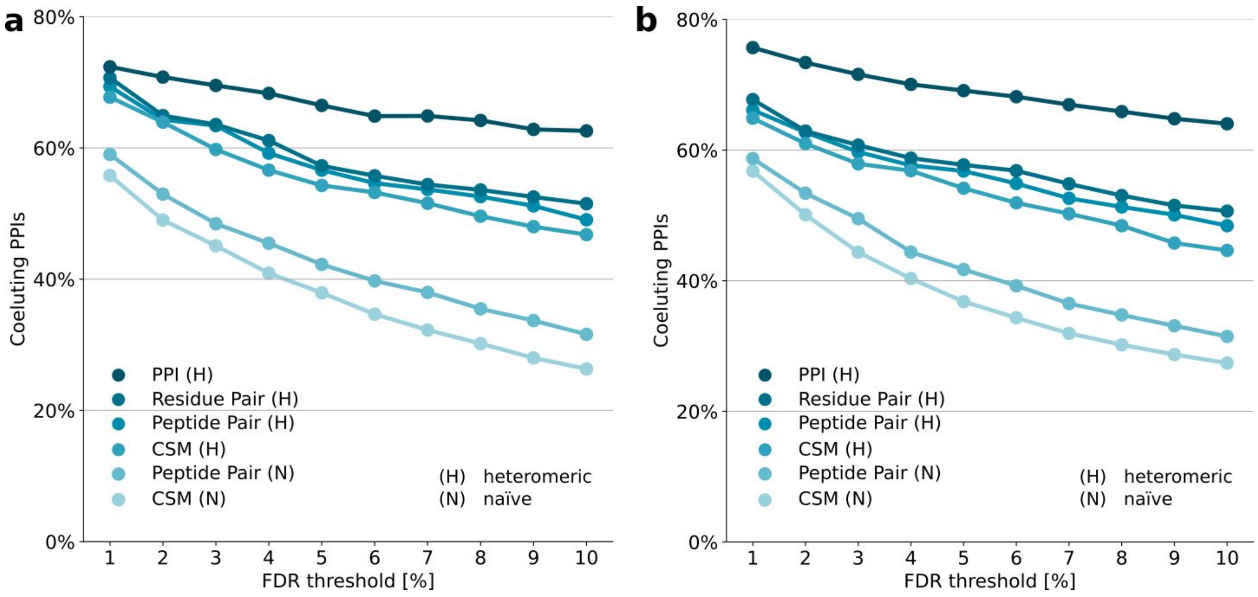


**Supplementary Figure 7: Results of positive control, coeluting PPIs.** Fraction of coeluting protein pairs (correlation coefficient > 0.5) among the heteromeric PPIs passing a given FDR threshold, applying different published FDR approaches separated for a) BS3 and b) DSSO.

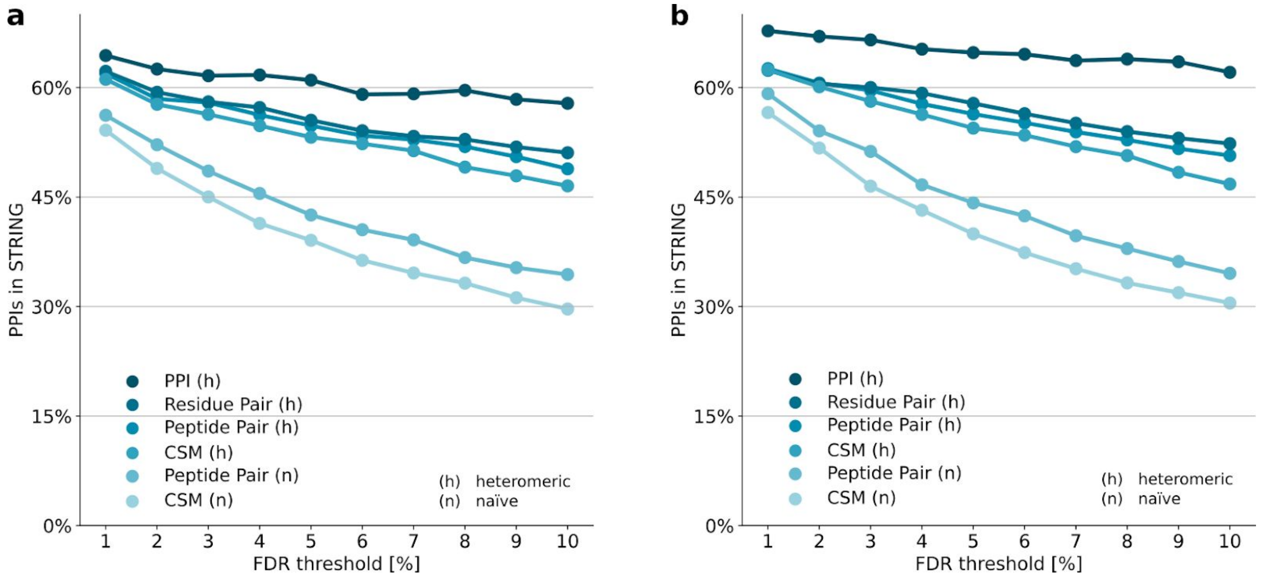


**Supplementary Figure 8: Results of alternative positive control, STRING database.** Fraction of protein pairs present in the STRING database (STRING combined score >= 150) among the PPIs passing a given FDR threshold, applying different published FDR approaches (see **Tab. S1**) for a) BS3 and b) DSSO.

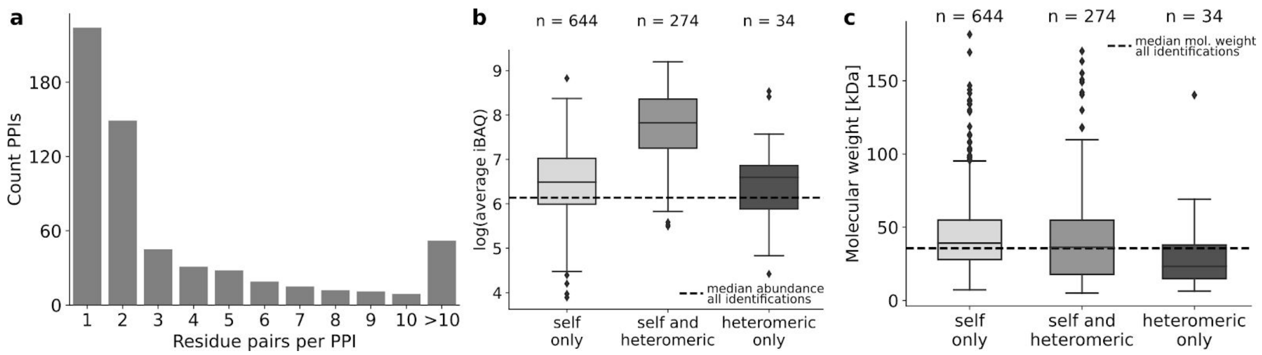


**Supplementary Figure 9: Properties of final crosslink PPI network.** a) Distribution of residue pairs per PPI. b) Abundance of identified crosslinked proteins in the respective categories (see **Fig. S5b**). “Heteromeric only” proteins are significantly more abundant than all identified proteins ($p < 0.05$ using a one-sided Kolmogorov–Smirnov test). c) “Heteromeric only” proteins tend to be smaller than the median of proteins in the database, and are therefore less likely to produce self-links.

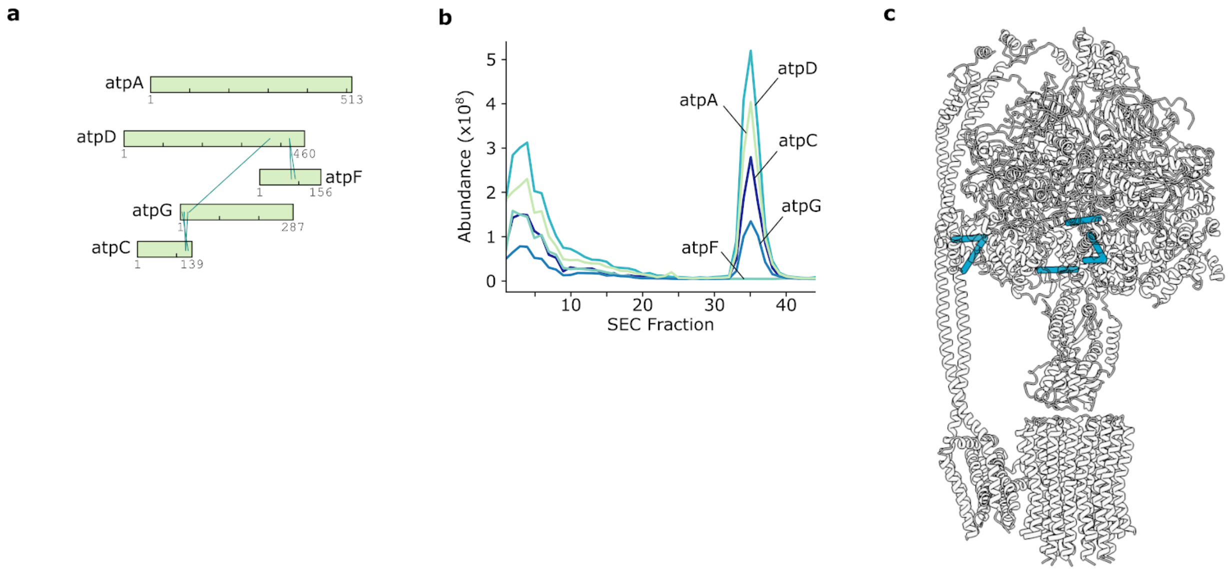


**Supplementary Figure 10: ATP synthase CLMS subnetwork.** a) PPI subnetwork of ATP synthase in xiVIEW[26]. Green shade on protein sequences illustrate sequence areas covered by the PDB model (PDB 5T4O). b) SEC-Coelution traces of ATP synthase subunits. Note that all subunits also coelute in a very early fraction, probably containing lipid vesicles. c) Structural model of ATP synthase with mapped heteromeric crosslinks (PDB 5T4O). All protein chains are colored in grey. Heteromeric links are below 35 Å and are colored in blue.

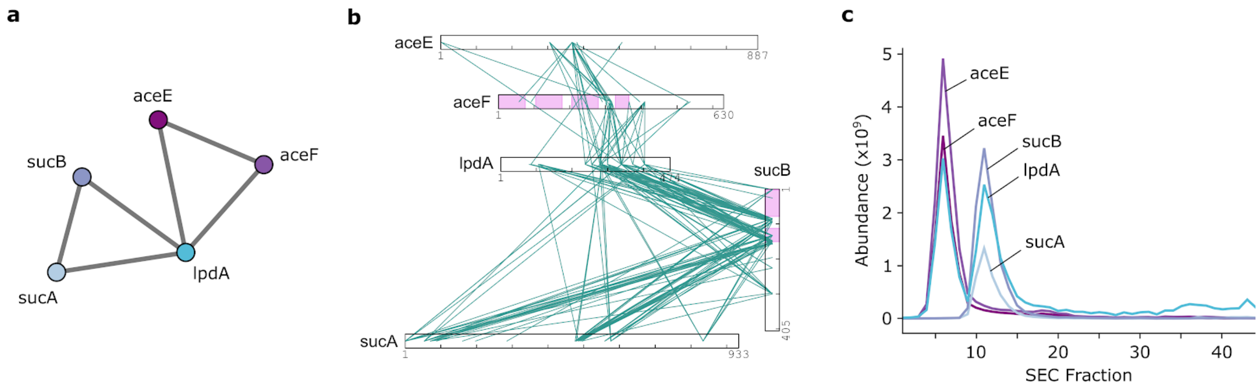


**Supplementary Figure 11: Pyruvate dehydrogenase CLMS subnetwork.** a) PPI subnetwork of pyruvate dehydrogenase and 2-oxoglutarate complexes with collapsed protein nodes and (b) with proteins shown as bars in xiVIEW[26]. c) Coelution of pyruvate dehydrogenase / 2-oxoglutarate complex components showing lpdA eluting with both.

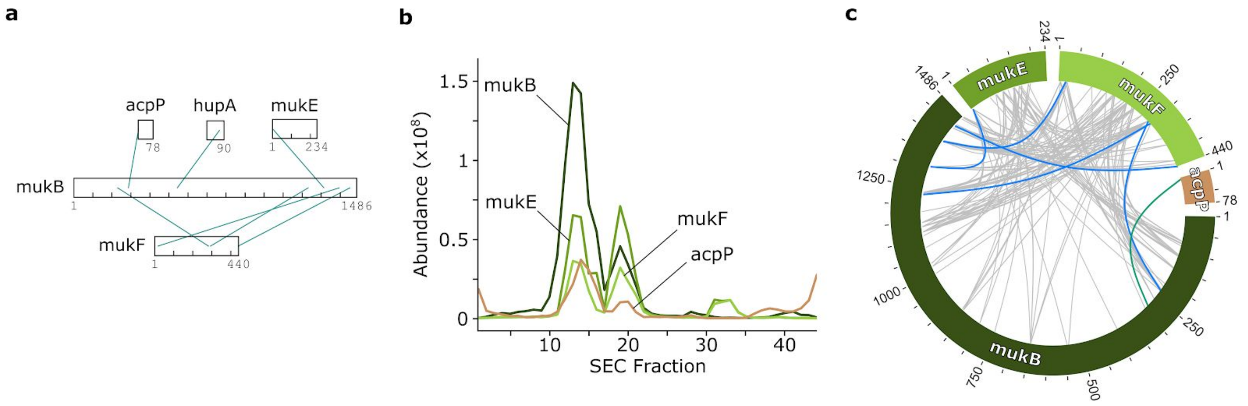


**Supplementary Figure 12: MukBEF complex CLMS subnetwork.** a) PPI subnetwork of MukBEF complex with proteins shown as bars in xiVIEW[26]. b) Coelution traces of MukBEF complex with its binder AcpP (acpP). The MukBEF complex (fraction 13) dissociated into another MukBEF assembly with lower occupancy for MukB (fraction 19) as judged by coelution. The interactor AcpP is a known binder of MukBEF that could be confirmed by CLMS and coelution. c) Crosslink distribution on MukBEF protein sequences compared to a recent *in-vitro* study[39]. Crosslinks are colored in blue if shared between the studies, in grey for *in-vitro* data only and in green if unique to this study.

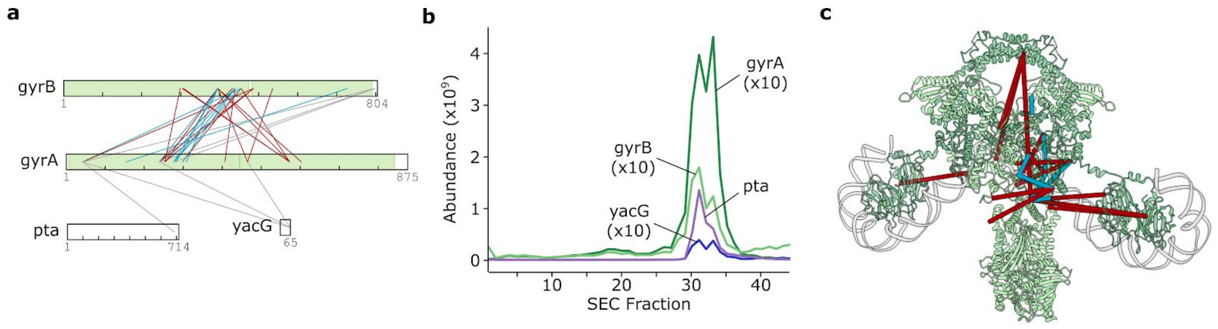


**Supplementary Figure 13: DNA gyrase CLMS subnetwork.** a) PPI subnetwork of DNA gyrase in xiVIEW[26]. Green shade on protein sequences illustrate sequence areas covered by the PDB model (PDB 6RKW). Heteromeric links are colored in blue if below 35 Å, red when exceeding and grey when absent from the model used. b) Coelution of DNA gyrase complex components with its inhibitor YacG and phosphate acetyltransferase Pta. The abundances for GyrA, GyrB and YacG were magnified 10-fold. c) Structural model of DNA gyrase with mapped heteromeric crosslinks (PDB 6RKW). GyrA is shown in dark green and GyrB in light green. DNA is shown in grey. For crosslink coloration see panel a. Of note, the DNA gyrase inhibitor YacG was suggested to associate with proteins involved in coenzyme A metabolism such as Pta[40], which is supported by this analysis.

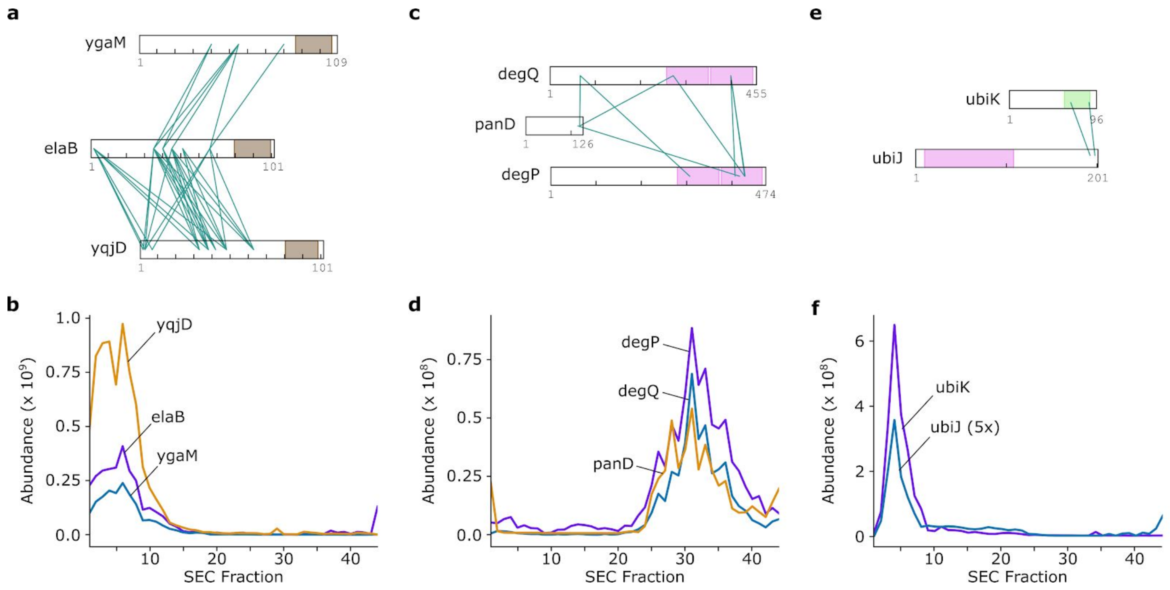


**Supplementary Figure 14: Novel PPIs identified.** Selected heteromeric PPI networks with proteins shown as bars in xiVIEW[26] (a, c, e) with their corresponding SEC elution traces (b, d, f). Coloration in panels a / c / e represent: transmembrane domains for elaB network; PDZ domains for degP and deg Q (pink); coiled-coil domain (green) for ubiK and SCP2 domain for ubiJ (pink). The abundance for ubiJ was magnified 5-fold.

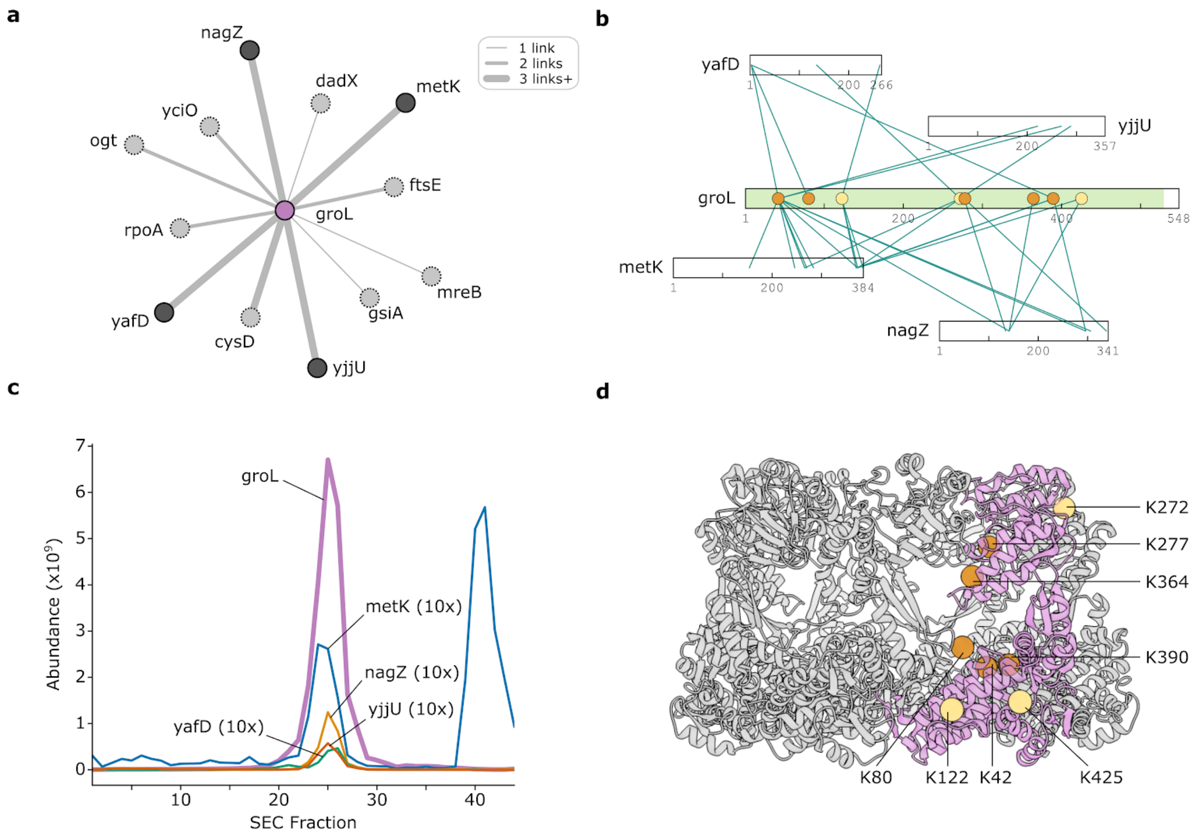


**Supplementary Figure 15: GroEL CLMS subnetwork.** a) PPI subnetwork of the chaperonin GroEL. High-confidence binders with darker nodes were selected for the following panels. b) Selected binders of GroEL shown as bars in xiVIEW[26]. Green areas mark GroEL's aligned region with a structural model (PDB 4PKO). Lysines on the inside of GroEL are highlighted with an orange circle, the ones on the outside with a circle in light yellow. c) Coelution of GroEL and its binders. The abundances of interactors were magnified 10-fold. d) GroEL structural model (PDB 4PKO) with crosslinked residues highlighted. One ring of the GroEL barrel assembly is shown with one subunit of GroEL colored in pink. Crosslinked lysine residues are indicated as spheres and colored as in panel b.

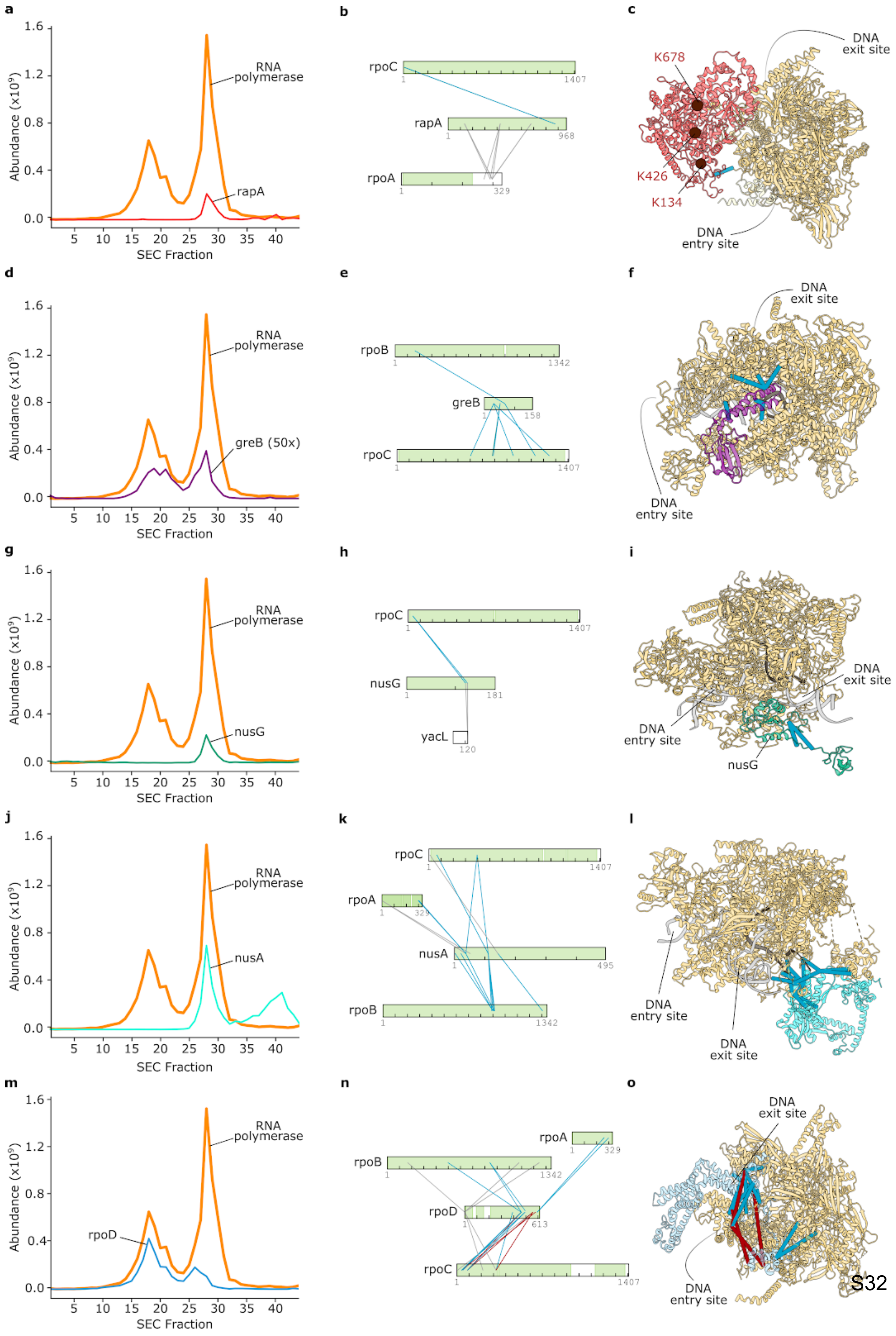
a
b
c
rpoC
rapA
rpoA
K678
K426
K134
DNA exit site
DNA entry site
d
e
f
rpoB
greB
greB (50x)
g
h
i
nusG
yacL
j
k
l
nusA
m
n
o
rpoD
RNA polymerase
Abundance (x10⁹)
SEC Fraction

**Supplementary Figure 16: Selected RNA polymerase binders.** Collected experimental support for heteromeric PPIs between RNAP and selected binders - rapA, greB, nusG, nusA and rpoD - is shown from SEC elution profiles (a, d, g, j, m), CLMS-based PPI screening in xiVIEW[26] (b, e, h, k, n) and matching to existing structural PPI models (c, f, i, l, o). The abundance of greB was magnified 50-fold. Green shade on protein sequence bars illustrate areas covered by PDB models that were used to map heteromeric links onto structures. In protein crosslink diagrams and protein structural models, heteromeric crosslinks satisfying euclidean distance thresholds of 35 Å are shown in blue while longer links are colored in red. The structural models used were: rapA (PDB 4S20), greB (PDB 6RIN), nusG (PDB 5MS0), nusA (PDB 6FLQ) and rpoD (PDB 4ZH3). If present in the model, DNA is colored in grey and RNA in black. Labeled spheres in panel c mark lysine residues crosslinked to C-termini of RNAP's α-subunits (absent from this model).

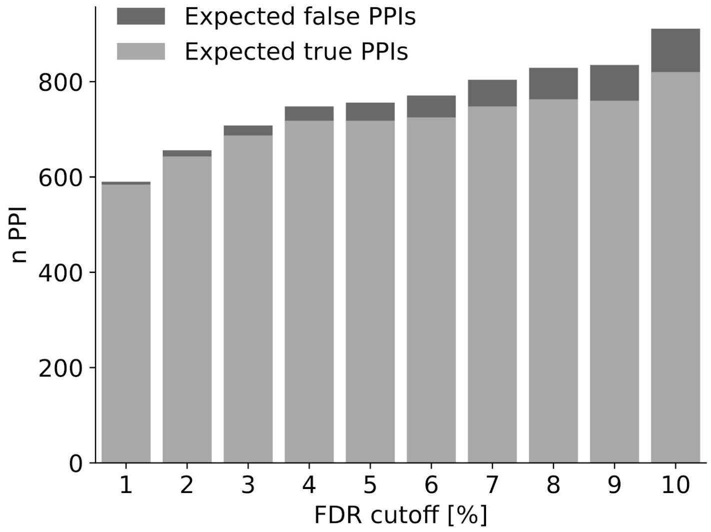


**Supplementary Figure 17: PPI numbers at different FDR thresholds.** The overall heteromeric PPI counts and the respective fraction of false PPIs are shown for varying FDR thresholds. Expected false PPIs were calculated based on decoy matches. For this dataset increasing FDR thresholds up to 4% add more true than false matches.
